# A CRISPR-Based Humanized Model Reveals Cooperative Role of STAG2 Loss in Familial GATA2-Deficient MDS Progression

**DOI:** 10.64898/2026.01.30.702879

**Authors:** Grace Freed, Miguel Quijada-Álamo, Linda Lee, Nikita Podar, Subrina Autar, Saul Carcamo, Persephone Fiore, Ke Wang, Isabella G. Martinez, Mimi Zhang, Shayan Saniei, Clifford Chao, Levan Mekerishvili, Zayna Diaz, Sai Ma, Dan Hasson, Elvin Wagenblast

**Affiliations:** Department of Oncological Sciences, Icahn School of Medicine at Mount Sinai, New York, NY, USA; Tisch Cancer Institute, Icahn School of Medicine at Mount Sinai, New York, NY, USA; Black Family Stem Cell Institute, Icahn School of Medicine at Mount Sinai, New York, NY, USA; Center for Advancement of Blood Cancer Therapies, Icahn School of Medicine at Mount Sinai, New York, NY, USA; Tisch Cancer Institute Bioinformatics for Next Generation Sequencing (BiNGS) core, Icahn School of Medicine at Mount Sinai, New York, NY, 10029, USA; Division of Hematology and Medical Oncology, Department of Medicine and Meyer Cancer Center, Weill Cornell Medicine, New York, NY, USA; Sandra and Edward Meyer Cancer Center, Weill Cornell Medicine, New York, NY, USA; Physiology, Biophysics and Systems Biology Graduate Program, Weill Cornell Medicine, New York, NY, USA; New York Genome Center, New York, NY, USA; Department of Genetics and Genomic Sciences, Icahn School of Medicine at Mount Sinai, New York, NY 10029, USA; Institute of Genomic Health, Icahn School of Medicine at Mount Sinai, New York, NY 10029, USA; Skin Biology and Diseases Resource-based Center, Icahn School of Medicine at Mount Sinai, NYC, NY 10029, USA; Department of Pediatrics, Division of Pediatric Hematology-Oncology, Icahn School of Medicine at Mount Sinai, New York, NY, USA; Mindich Child Health & Development Institute, Icahn School of Medicine at Mount Sinai, New York, NY, USA

## Abstract

Myelodysplastic syndrome (MDS) is a heterogeneous myeloid malignancy driven by hematopoietic stem cell dysfunction, leading to ineffective hematopoiesis and cytopenias. Familial GATA2 deficiency is the most common cause of Myelodysplastic syndrome in adolescents, with progression often accelerated by co-occurring mutations, notably STAG2 loss-of-function. Using CRISPR/Cas9-mediated genome engineering in primary human fetal liver-derived hematopoietic stem cells and xenotransplantation in mice, we modeled GATA2-deficient Myelodysplastic syndrome with acquired STAG2 loss to investigate disease initiation and progression. While GATA2 deficiency alone had minimal short-term impact in our model, combined GATA2 and STAG2 loss increased hematopoietic stem cell maintenance and self-renewal, induced a myeloid-lineage bias, and expanded primitive progenitors. Single-cell transcriptional profiling revealed upregulation of stemness genes and inflammatory pathways. This humanized model faithfully recapitulates high-risk GATA2-deficient Myelodysplastic syndrome, providing mechanistic insight into how cooperative mutations drive stem cell expansion, inflammatory signaling, and myeloid skewing.

**Visual Abstract:** 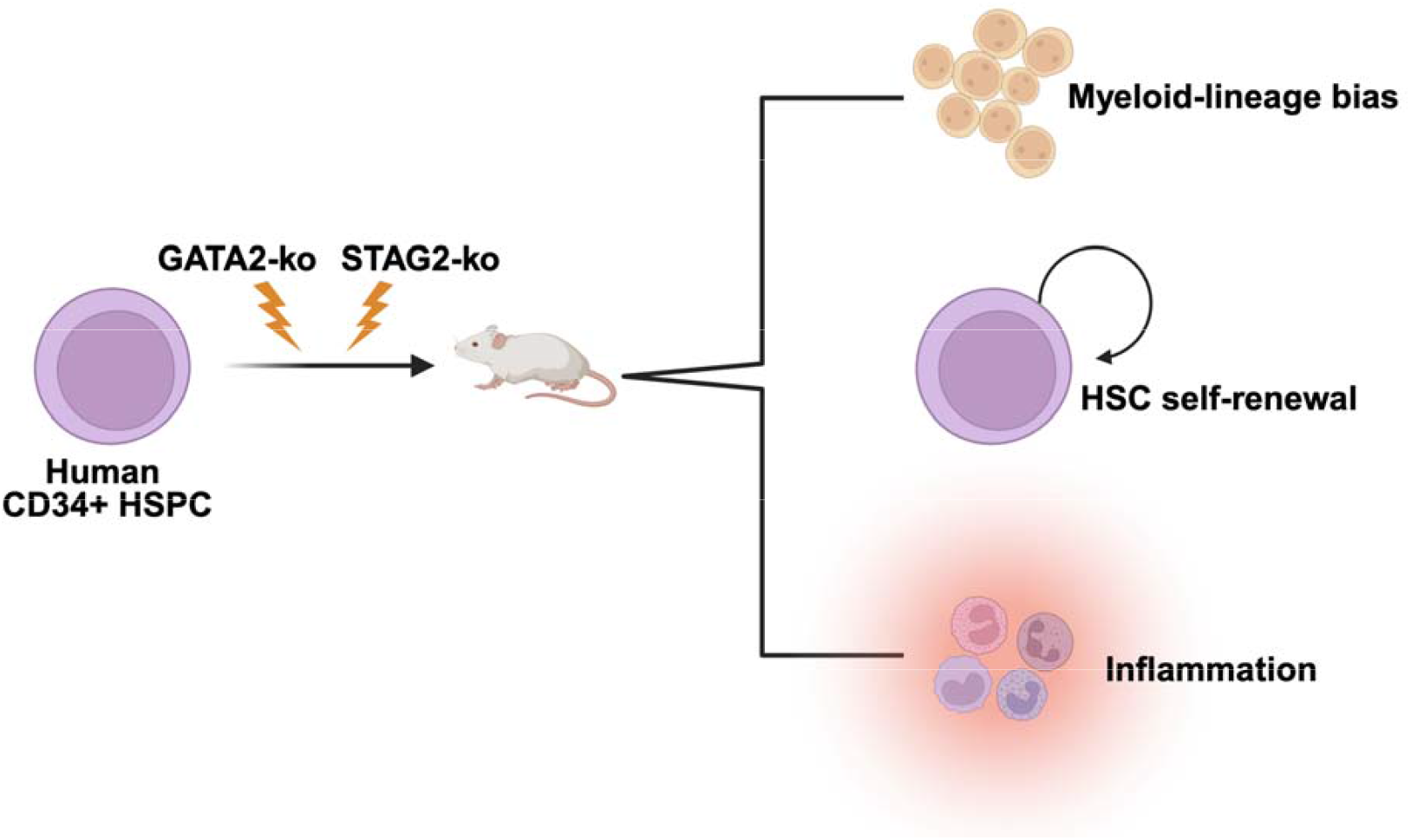

**Key Points:**

- Humanized model of familial GATA2-deficiency requires the loss of STAG2 for progression to an MDS disease phenotype
- GATA2-ko+STAG2-ko increase HSC self-renewal, induce a myeloid-lineage bias, and trigger an inflammatory transcriptional program

## Introduction

Myelodysplastic syndrome (MDS) comprises a heterogeneous group of myeloid malignancies arising from a diverse landscape of mutations in hematopoietic stem cells (HSCs). It is characterized by the accumulation of immature or dysfunctional blood cells due to impaired HSC function, deregulated apoptosis, and dysregulation of inflammatory and innate immune pathways^1^. Clinically, MDS presents with anemia, increased susceptibility to infection, and bleeding, and progresses to acute myeloid leukemia (AML) in ∼30-40% of individuals^2^. A hallmark of MDS pathogenesis is heightened inflammatory signaling^3^. Inflammation promotes clonal expansion of malignant HSCs in MDS, leading to an expansion of these stem and progenitor cell compartments^4^. However, the role of inflammation in the initiation of MDS, and how it could potentially be therapeutically exploited, is not fully understood.

While MDS predominantly affects adults over the age of 60, certain genetic predispositions, such as familial GATA2 deficiency, are strongly associated with pediatric-onset disease^5^. Familial GATA2 deficiency, caused by heterozygous germline mutations in the *GATA2* gene, results in a spectrum of immunodeficiencies and myelodysplasias^6^ and is the most common genetic cause of MDS in adolescents^7^. While GATA2 deficiency does not confer a worse prognosis to MDS patients, individuals with GATA2 deficient MDS are at a high risk for progressing to AML^5^, underpinning the importance of further investigation.

GATA2 encodes a master regulator of hematopoiesis and a member of the GATA transcription factor family. The protein contains two zinc-finger domains (ZF1 and ZF2) that bind GATA motifs across the genome: ZF1 primarily mediates protein-protein interactions, whereas ZF2 facilitates transcription through protein-DNA binding^7^. In postnatal hematopoiesis, GATA2 is highly expressed in HSCs and hematopoietic progenitors, promoting myeloid-erythroid lineage development^7^. The mutational landscape of GATA2 deficiency is diverse, consisting of missense mutations mostly located in the zinc finger domains and null mutations located upstream of the zinc finger domains^8^. Genetically engineered mouse models have shown that GATA2 deficiency impairs HSC self-renewal, maintenance, and survival and promotes myeloid lineage skewing in mice^9,10^. A recent study further demonstrated impaired HSC self-renewal and survival using a humanized model of GATA2 deficiency^11^. In a study of a large patient cohort, it was observed that patients with GATA2 missense mutations have a high-risk of developing leukemia^12^. Interestingly, another study of individuals harboring germline GATA2-deficiency show that some individuals remain asymptomatic, even with mutations in GATA2, but can progress to a malignant state upon acquisition of somatic mutations^10^. These findings motivated us to interrogate the functional role of GATA2 deficiency in the initiation of MDS using a humanized model, and whether GATA2 deficiency on its own is sufficient to produce a phenotype.

Familial GATA2-deficient MDS is frequently accompanied by additional somatic mutations that drive progression to high-risk MDS or transformation to AML^13^. Commonly co-occurring mutations include *STAG2, ASXL1*, and *SETBP1*, each associated with worse prognosis^14^. *STAG2* encodes a core component of the cohesin complex, which regulates chromatin architecture during cell division, transcription, and DNA repair. Mutations in *STAG2* account for over half of all cohesin mutations in myeloid malignancies and occur in ∼25% of GATA2-deficient MDS cases^15,16^. In a genetically engineered mouse model, STAG2 loss alone increases HSC and progenitor self-renewal and induces myeloid-biased differentiation^17^. Clinically, GATA2-deficient MDS patients with *STAG2* mutations are categorized as high-risk and have an increased chance of AML progression; however, this cooperative disease progression has not been functionally characterized or faithfully modeled in human HSCs.

In this study, we set out to generate a humanized model of GATA2-deficient MDS encompassing one of the most frequent co-occurring somatic mutations, STAG2 loss-of-function. Using CRISPR/Cas9-mediated genome engineering, we knocked out GATA2 and STAG2 individually and in combination in primary human fetal liver-derived CD34+ hematopoietic stem and progenitor cells (HSPCs), which was followed by xenotransplantation into immunodeficient mice. We used fetal liver-derived HSPCs in this study to recapitulate the fetal context of a germline GATA2 mutation. These models allowed us to investigate the progression of familial GATA2-deficient MDS to high-risk disease following STAG2 loss.

## Methods

### Human patient samples

Human fetal liver samples were obtained from elective pregnancy terminations from the Developmental Origins of Health and Disease (DOHaD) Biorepository at the Icahn School of Medicine at Mount Sinai with written informed consent following guidelines approved by the Icahn School of Medicine at Mount Sinai Institutional Review Board. Fetal liver samples were collected between 16 to 21 weeks’ gestation.

### Animal studies

All mouse experiments were approved by the Icahn School of Medicine at Mount Sinai Institutional Animal Care and Use Committee. We confirm that all experiments conform to the relevant regulatory and ethical standards. All xenotransplantations were performed in 8 to 12-week-old *NOD*.*Cg-Prkdc*^*scid*^*Il2rg*^*tm1Wjl*^*/SzJ* (NSG) mice (JAX) that were sublethally irradiated with 230 cGy, 24 hours before transplantation. Human cells were transplanted via intrafemoral injections.

## Results

### Combined GATA2 and STAG2 Loss Induces Myeloid-Lineage Bias *In Vivo*

To investigate the impact of GATA2 deficiency (GATA2-ko) and STAG2 loss (STAG2-ko) individually and in combination, we established a humanized *in vivo* model using primary human fetal liver (FL)-derived CD34+ HSPCs. Cells were CRISPR/Cas9-edited resulting in control (targeting the olfactory receptor OR2W5), GATA2-ko, STAG2-ko, and GATA2-ko + STAG2-ko and xenotransplanted through intra-femoral injections into immunodeficient NSG mice, with human engraftment in the bone marrow analyzed after 12 weeks (**Figure 1A**). CRISPR/Cas9 editing was accomplished by generating custom guide RNAs (gRNAs) targeting each gene, followed by binding of the gRNAs to Cas9 protein, and electroporation of the ribonucleoprotein complex into primary HSPCs. GATA2 editing targeted exon 6 on chromosome 3, resulting in a loss-of-function missense mutation in the zinc-finger 2 DNA-binding domain (**Figure 1B**). Our gRNA editing resulted in cells with most alleles carrying a +1 bp insertion (**Figure 1C**). To confirm the heterozygous state of the GATA2 mutation, consistent with patient genotypes, we cultured GATA2-edited CD34+ FL cells in methylcellulose colony-forming media. After 12 days, single cell-derived colonies were isolated for DNA extraction and genotyped. Sanger sequencing-based editing efficiency indicated 70% were heterozygous and 30% were homozygous for GATA2-ko (**Figure 1D**). These results demonstrate that our CRISPR/Cas9-based editing strategy generated a mix of heterozygous and homozygous GATA2-deficient cells. For STAG2 knockout editing, we used a previously validated gRNA targeting exon 11 on chromosome X, resulting in a complete loss-of-function^18^.

**Figure 1.**
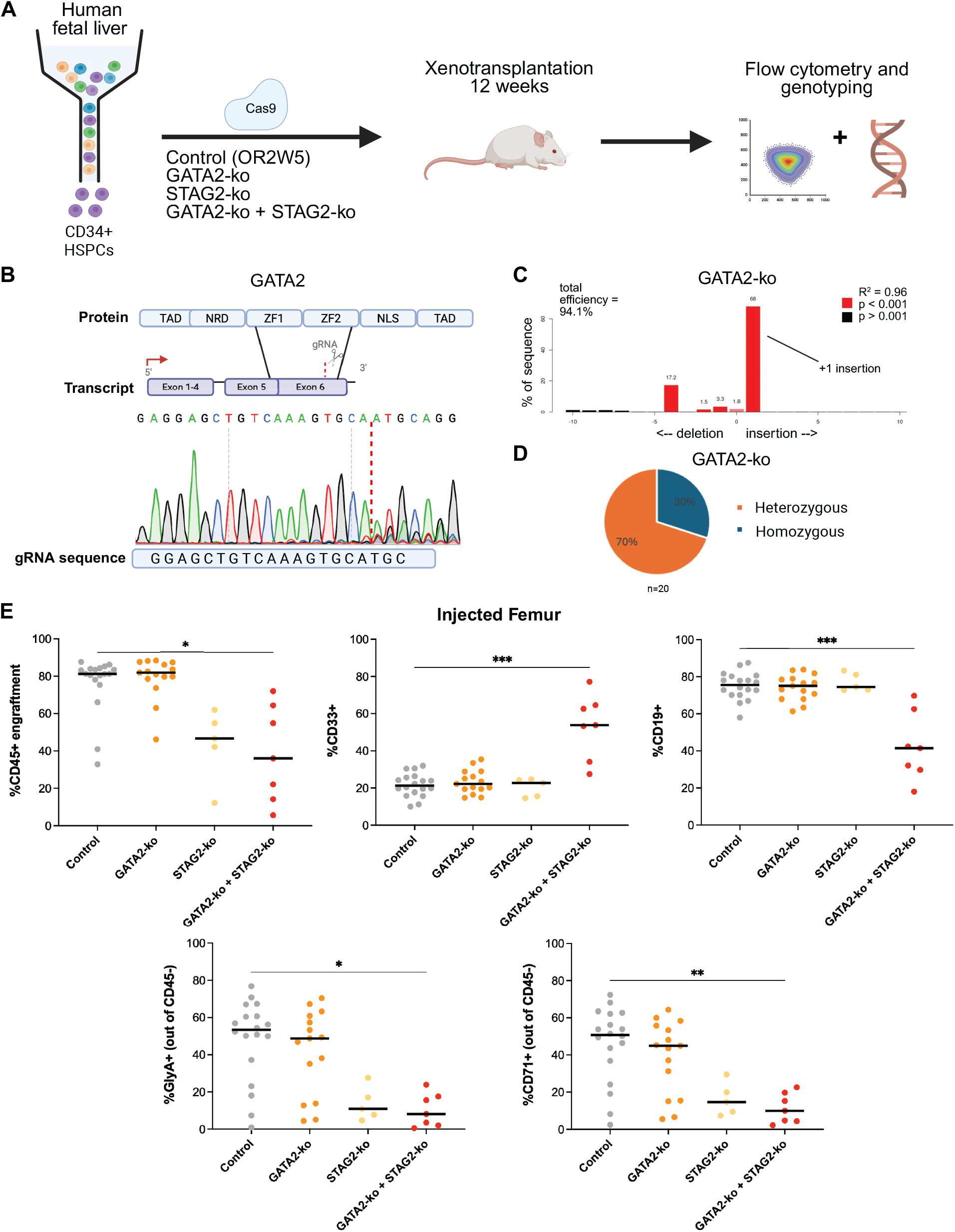
Humanized xenograft models of GATA2 deficient MDS with STAG2 knockout shows myeloid-lineage bias *in vivo*. (A) Experimental overview of fetal liver-derived CD34+ HSPC isolation and CRISPR/Cas9-mediated engineering followed by intrafemoral injection into NSG mice, allowing engraftment for 12 weeks before immunophenotyping. (B) Experimental design of developing a GATA2 gRNA to induce a double-stranded break in exon 6 of the gene, resulting in loss of function in the ZF2 domain. (C) Example TIDE results showing overall efficiency and the most frequent indels resulting from GATA2 gene editing. (D) Results from Sanger sequencing of colony-forming assay, showing genotyping results of single cell-derived colonies and categorized as either heterozygous or homozygous, n = 20 colonies. (D) Immunophenotypic analysis of the injected bone marrow of control, GATA2-ko, STAG2-ko, and GATA2-ko+STAG2-ko primary xenografts. CD45: overall human cell engraftment; CD33: myeloid lineage; CD19: B-cell lymphoid lineage; CD71: immature erythroid marker; GlyA: mature erythroid marker. The line represents the mean value. *p < 0.05, **p < 0.01, ***p < 0.001, and ****p-value <0.0001, Kruskal-Wallis Test, n=5-18 mice each.

To further validate GATA2 deficiency in our primary CRISPR/Cas9 engineered HSPCs, we employed single-cell genotyping of targeted loci with chromatin accessibility (GoT-ChA)^19^ upon GATA2-ko in FL-derived CD34+ HSPCs and 12 weeks of *in vivo* engraftment. GoT-ChA allowed us to simultaneously profile the chromatin accessibility landscape and GATA2 genotype at single-cell resolution, thereby distinguishing the epigenetic landscape of mutant from wild-type cells within the same sample. The UMAP generated from this dataset showed a mix of GATA2 homozygous mutant, heterozygous mutant, and wild-type cells in our sample (**Supplementary Figure 1A**). We obtained a high genotyping efficiency of 52% with 4,355 genotyped cells. (**Supplementary Figure 1B**). Our data shows 12% heterozygous GATA2-ko, 63% homozygous GATA2-ko, and 25% wild-type cells. The percentages of heterozygous and homozygous GATA2-ko cells here differ from our *in vitro* colony data with a higher number of homozygous mutant cells. This could be due to different clonal selection during *in vivo* engraftment compared to the *in vitro* colony assay, resulting in the selection of different clones over time. Importantly, we found increased chromatin accessibility of GATA family transcription factor motifs upon GATA2-ko, specifically in the cycling progenitors (**Supplementary Figure 1C**). Therefore, GoT-ChA allowed us to validate that GATA2 loss-of-function has a direct consequence on open chromatin accessibility of GATA motifs, potentially indicating a feedback loop.

In order to assess the effects of GATA2 and STAG2 loss-of-function *in vivo*, we conducted immunophenotypic analysis of the bone marrow of the xenografted mice, both in the injected and non-injected femur. In the injected bone marrow of FL xenografted mice, STAG2-ko alone and GATA2-ko + STAG2-ko human grafts showed a significant decrease in overall human CD45+ engraftment. Interestingly, GATA2-ko + STAG2-ko grafts showed a marked shift in lineage composition, with a significant 2.5-fold increase in CD33+ myeloid cells and a corresponding significant decrease in CD19+ B-lymphoid cells (**Figure 1D)**. Importantly, neither GATA2-ko nor STAG2-ko alone altered lineage distribution within the myeloid and lymphoid lineages at 12 weeks post-engraftment. Finally, GATA2-ko + STAG2-ko grafts showed a significantly decreased CD71+ immature erythroid and GlyA+ mature erythroid output in the mouse xenografts (**Figure 1D**). Similar results were seen in the non-injected bone marrow of the xenografted mice (**Supplementary Figure 1D**). Overall, these immunophenotyping results indicate a myeloid-lineage bias in GATA2-ko + STAG2-ko xenografts due to a cooperative role between these two genes, as they are not seen when these genes are individually knocked out.

### Single-Cell Profiling Reveals Expansion of Primitive Hematopoietic Cells upon GATA2/STAG2 Loss

Based on the results from the immunophenotypic analysis, we wanted to further characterize the different cell types present in each xenograft model, specifically the stem and progenitor compartment, which requires a high resolution to capture. Therefore, we performed single-cell RNA sequencing (scRNA-seq) on CD45+ cells from our FL-derived xenograft models, enriched with 20% CD34+CD19-primitive HSPCs to map the transcriptional landscape of control, GATA2-ko, STAG2-ko and GATA2-ko + STAG2-ko grafts (**Figure 2A**). Using a single-cell reference dataset^20^, we annotated hematopoietic lineages in our merged scRNA-seq atlas. A total of ∼45,000 high-quality cells were analyzed, revealing 23 transcriptionally distinct clusters ranging from HSCs to mature myeloid and lymphoid lineages (**Figure 2B**). We generated a merged UMAP of the ∼45,000 cells from all xenograft conditions combined, showing distinct clusters of single cells based on gene expression data from each cell. Further, UMAPs representing each individual xenograft condition were generated (**Supplementary Figure 2A**). Using the cell type annotations in our individual UMAPs, the proportion of progenitor cell types out of the CD34+CD19-fraction in each xenograft condition was determined. Notably, GATA2-ko + STAG2-ko grafts showed a marked expansion of primitive populations, including HSCs, cycling progenitors, and early granulocyte-monocyte progenitors (early GMPs) compared to control, GATA2-ko, and STAG2-ko grafts (**Figure 2C**). An expansion of progenitor and immature cells in the bone marrow is a hallmark of the MDS disease state^1^, especially with a STAG2 mutation^17^.

**Figure 2.**
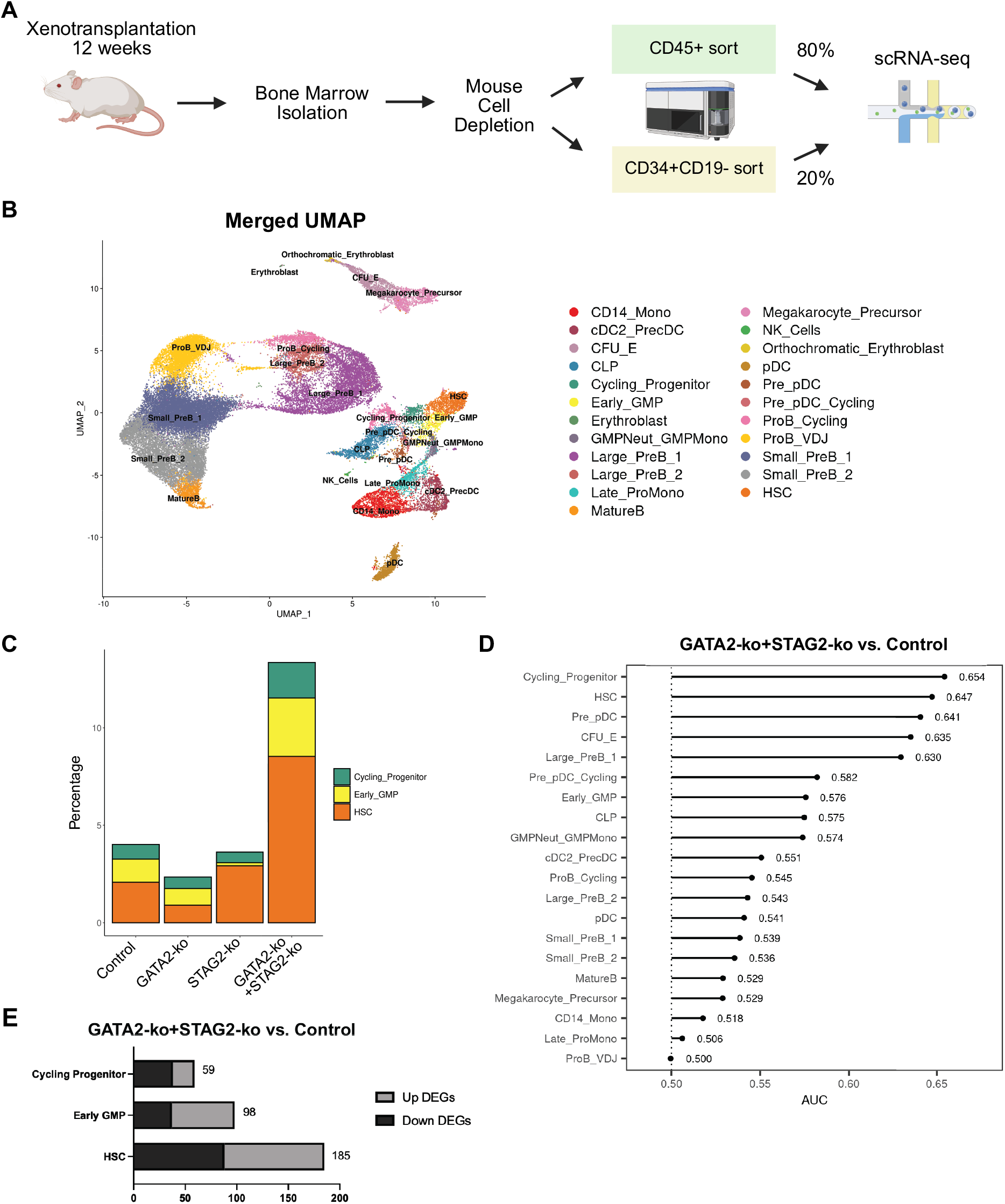
Single-cell transcriptional analysis of GATA2-ko + STAG2-ko xenografts shows marked expansion of stem and progenitor cells. (A) Experimental overview of primary xenograft mouse cell depletion and sorting strategy for scRNA-seq, resulting in 80% CD45+ human cells with 20% CD34+CD19-HSPC enrichment. (B) UMAP of merged primary xenografts including control, GATA2-ko, STAG2-ko, and GATA2-ko + STAG2-ko with cell-type annotations. (C) Proportions of HSCs, early GMPs, and cycling progenitors in each condition, shown as a percentage of total cells. (D) Result from Augur bioinformatics analysis showing prioritization of cell types based on their molecular response to perturbation. Showing comparison between GATA2-ko + STAG2-ko versus matched control. (E) Table showing upregulated and downregulated differentially expressed genes of the highest priority cell types, as assigned by Augur, between GATA2-ko + STAG2-ko versus control.

We next applied Augur^21^, a computational tool that ranks cell types by the magnitude of their transcriptional response to perturbation, to our single-cell dataset. This approach is particularly useful for assessing the contribution of rare or underrepresented cell types to a phenotype, as it avoids the bias of differential expression analyses toward more abundant populations. We used Augur to determine whether the expansion of primitive cell types observed in GATA2-ko + STAG2-ko xenografts is accompanied by a strong, cell-intrinsic transcriptional response to the double knockout mutation. This analysis revealed that cycling progenitors and HSCs showed the greatest transcriptional response in the GATA2-ko + STAG2-ko condition relative to their matched control, confirming our findings (**Figure 2D**). In parallel, differential gene expression analysis revealed a high number of differentially expressed genes in HSCs and early GMPs compared to control (**Figure 2E**). Interestingly, despite ranking highest in the Augur analysis, cycling progenitors showed fewer differentially expressed genes than these other progenitor subsets. The primitive populations, namely the HSCs and cycling progenitors, also showed a high degree of transcriptional response compared to their respective controls in the GATA2-ko and STAG2-ko single-mutant conditions (**Supplementary Figure 2B and 2C**). However, they were not the most transcriptionally perturbed populations in either single-mutant condition alone, whereas they are the top-ranked populations in the combined GATA2-ko + STAG2-ko condition. This indicates that while each individual gene disruption contributes partially to the transcriptional perturbation and overall expansion of these cell types, the combination of both mutations is required to fully manifest the transcriptional phenotype we observed in the double mutant HSCs and cycling progenitors. In fact, these cell types are significantly more transcriptionally perturbed in the GATA2-ko + STAG2-ko condition when directly compared to the GATA2-ko condition alone (**Supplementary Figure 2D**). Given the pronounced expansion of the stem and progenitor compartment and the distinct transcriptional alterations in the GATA2-ko + STAG2-ko condition, we prioritized these populations for downstream analyses.

### GATA2/STAG2 Double Knockout HSCs Exhibit Inflammatory Signatures and TNF-α Sensitivity

To identify transcriptional pathways enriched in the HSC compartment of the GATA2-ko + STAG2-ko xenografts, we performed differential gene expression analysis between this population and the corresponding cell population in the control xenograft. The GATA2-ko + STAG2-ko HSCs showed upregulation of inflammatory programs, most notably TNF-α, interferon-γ, and interferon-α response pathways (**Figure 3A**). Furthermore, pathways related to *E2F* and *MYC* target genes were downregulated in these cells, suggesting a differentiation block and a resulting accumulation of GATA2-ko + STAG2-ko HSCs^22^. Similarly, we conducted differential gene expression analysis between the cycling progenitors and the early GMPs in the GATA2-ko + STAG2-ko xenografts and their corresponding cell types in the control xenograft. The cycling progenitors in the GATA2-ko + STAG2-ko condition showed an upregulation of inflammatory pathways, specifically the interferon-γ response pathway (**Supplementary Figure 3A**). The early GMPs exhibited a remarkably similar transcriptional program to the HSCs, with upregulation of TNF-α signaling response and downregulation of *MYC* target pathways (**Supplementary Figure 3B**). Notably, when comparing the HSC compartment in GATA2-ko alone and STAG2-ko alone versus in the control xenograft, this HSC self-renewal and inflammation phenotype was not observed (**Supplementary Figure 3C and 3D)**. In fact, pathways related to *MYC* target genes were upregulated in the GATA2-ko condition, indicating stem cell differentiation (**Supplementary Figure 3C**). Pathways related to *MTORC1* signaling were upregulated in the STAG2-ko condition, indicating the contribution of the STAG2-ko to an increase in cell growth and proliferation^23^ (**Supplementary Figure 3D**). Finally, comparison of the HSC compartment in the GATA2-ko + STAG2-ko to the HSC compartment in the GATA2-ko alone condition revealed the upregulation of inflammatory pathways and downregulation of *MYC* signaling pathways (**Supplementary Figure 3E**). This confirms that the transcriptional phenotype is significantly pronounced in and unique to the combination condition. A heatmap summarizing the different enriched pathways that are upregulated and downregulated in the HSC compartment across all comparisons of conditions is shown (**Figure 3B**).

**Figure 3.**
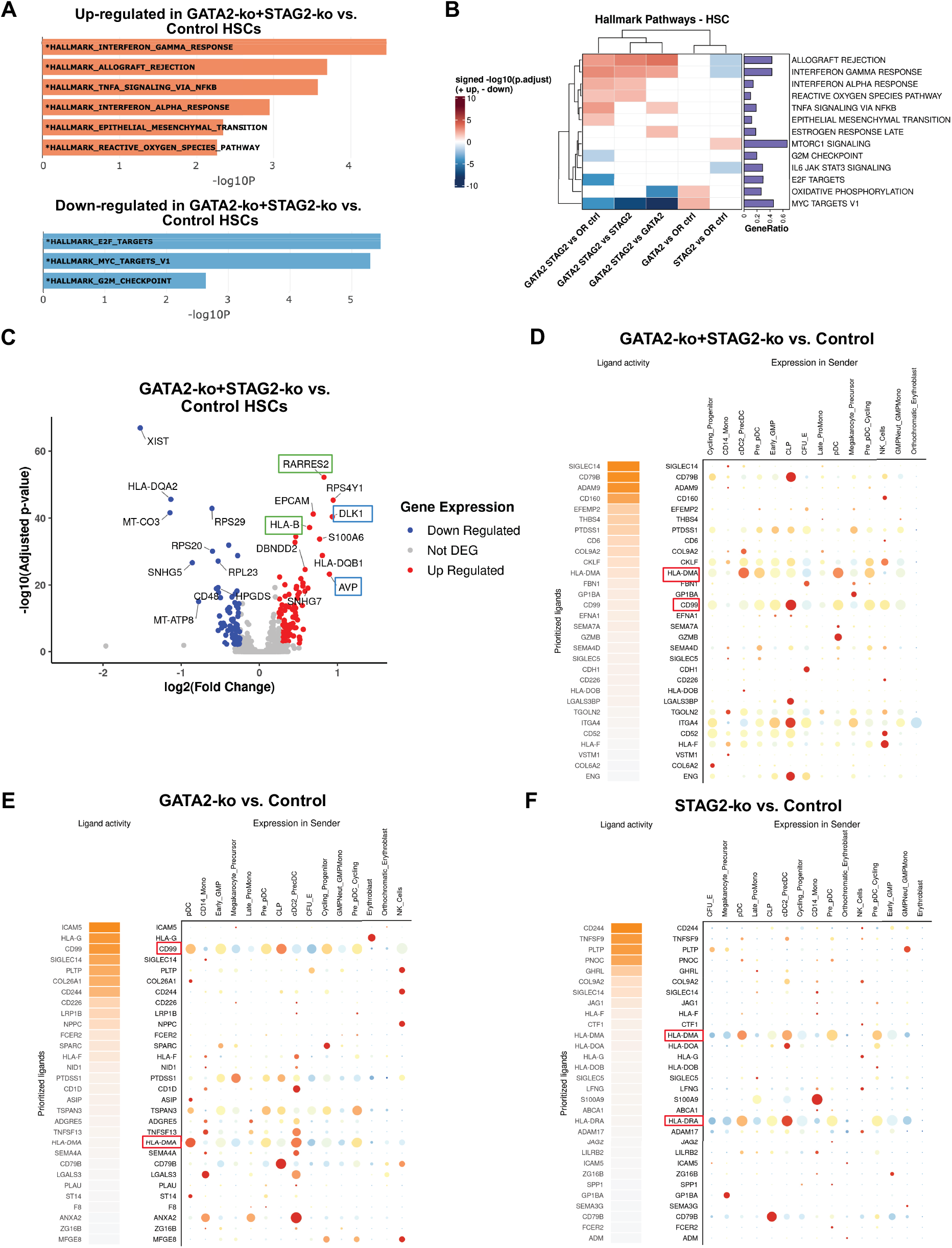
Single-cell RNA sequencing data reveals upregulation of HSC maintenance and inflammatory signaling in GATA2-ko + STAG2-ko xenografts. (A) Over Representation Analysis (ORA) on differentially expressed HSC genes between GATA2-ko + STAG2-ko versus control. Enriched pathways and biological processes were identified based on differentially expressed genes. Significant hallmark pathways are highlighted in bold and with an asterisk (*) based on p-value < 0.05. (B) Heat map of enriched hallmark pathways in the HSC compartment showing every comparison between conditions. Red indicates enrichment in upregulated genes and blue indicates enrichment in downregulated genes. Only significant hallmark pathways with a p-value < 0.05 are included. (C) Volcano plot of differentially expressed genes between GATA2-ko + STAG2-ko and control in the HSC compartment. Genes associated with inflammation are marked in green and genes associated with self-renewal are marked in blue. All genes with significant adjusted p-values < 0.05 and log2FC > 1 are colored. (D-F) Results from NicheNet computational analysis showing ligand-target gene interactions with HSCs as the receivers and all cells excluding lymphoid cells as the senders, comparing (C) GATA2-ko + STAG2-ko versus matched control, (D) GATA2-ko versus matched control, and (E) STAG2-ko versus matched control.

Several genes associated with stem cell quiescence and maintenance were significantly upregulated in GATA2-ko + STAG2-ko HSCs compared with control, including *DLK1* and *AVP* (**Figure 3B)**. *DLK1* is known to inhibit HSC differentiation^24^, and *AVP* is a marker of long-term HSCs that is also linked to stress-induced myeloid differentiation in HSCs^25^. This gene profile supports the observed expansion of HSCs in this condition. *DLK1* was also upregulated in the cycling progenitor population, indicating maintenance of self-renewal in these cells (**Supplementary Figure 3F**). In addition, the HSC compartment showed upregulation of inflammatory pathway genes, including *HLA-B* and *RARRES2* (**Figure 3B**). These same inflammatory genes were similarly upregulated in cycling progenitors and early GMPs in the GATA2-ko + STAG2-ko condition (**Supplementary Figure 3F and 3G**).

To identify ligand-target gene interactions between sender and receiver cell populations in our scRNA-seq dataset, we applied the computational method NicheNet^26^. This analysis revealed that MHC-II class genes, such as *HLA-DMA*, were upregulated in GATA2-ko + STAG2-ko HSCs primarily in response to signals from dendritic cell precursors (pre-cDCs and pDCs) (**Figure 3C**). These MHC-II genes are frequently upregulated in HSCs in inflammatory disease contexts, including MDS, and are associated with reduced HSC apoptosis and impaired differentiation^27^. *HLA-DMA* was also upregulated in GATA2-ko HSCs and STAG2-ko HSCs individually (**Figure 3D and 3E**). Furthermore, we observed an upregulation of *CD99* in both GATA2-ko + STAG2-ko HSCs and GATA2-ko HSCs primarily in response to signals from common lymphoid progenitors (CLPs) (**Figure 3C and 3D**). *CD99* upregulation has been described to promote self-renewal in HSCs in acute myeloid leukemias^28^. This *CD99* expression is notably absent in the STAG-2-ko HSCs and is prominent in GATA2-ko HSCs due to high ligand activity, suggesting that GATA2-ko is driving this phenotype in the combined knockout condition. On the other hand, the HLA gene over-expression is the more prominent phenotype in the STAG2-ko HSCs, suggesting that STAG2-ko plays a driving role in this phenotype in the combined condition (**Figure 3E**). Overall, these transcriptional analyses confirmed the necessity of the combined GATA2-ko + STAG2-ko condition to observe our MDS phenotype but elucidated that each gene knockout individually contributes unique and important elements to the combined phenotype.

### Combined GATA2 and STAG2 Loss Enhances Self-Renewal of Primitive Hematopoietic Cells

Given the expansion of HSCs and cycling progenitors in GATA2-ko + STAG2-ko xenografts observed by scRNA-seq, we assessed the *in vitro* self-renewal capacity of these cells. We focused on the GATA2-ko and GATA2-ko + STAG2-ko conditions for the following functional experiments, as these are the natural mutational conditions observed in patients with GATA2-deficient MDS. To assess self-renewal capacity, CD34+ FL cells edited for control, GATA2-ko and GATA2-ko + STAG2-ko were cultured in methylcellulose colony-forming media and serially replated three times to evaluate their ability to continuously generate colonies. At each passage, genotype-positive colonies were counted, and overall cell populations were analyzed by flow cytometry (**Figure 4A**). GATA2-ko + STAG2-ko cells produced more genotype-positive colonies after the secondary replating, relative to the number of colonies in the primary plating, compared to GATA2-ko and control. STAG2-ko cells also produced more genotype-positive colonies after the secondary replating, relative to the number of colonies in the primary plating. However, the increase of colonies in the STAG2-ko condition was noticeably less pronounced than that of GATA2-ko + STAG2-ko condition. In addition, GATA2-ko + STAG2-ko were the only condition to yield a high number of colonies after the tertiary replating (**Figure 4B**). Flow cytometry analysis of these colonies revealed an increase in the percentages of myeloid lineages in the GATA2-ko + STAG2-ko colonies and the STAG2-ko colonies over the course of the assay, with increased CD33+ myeloid cells and CD33+CD34+ immature myeloid cells (**Figure 4C**). Further, there was an overall reduction in the percentage of CD33+CD66b+ mature granulocytes in GATA2-ko + STAG2-ko colonies compared to the colonies that are produced in GATA2-ko, STAG2-ko, and control. While STAG2-ko alone promotes an immature myeloid lineage *in vitro* and slightly increases HSC self-renewal, the combined GATA2 and STAG2 loss is required for a pronounced effect on HSC self-renewal.

**Figure 4.**
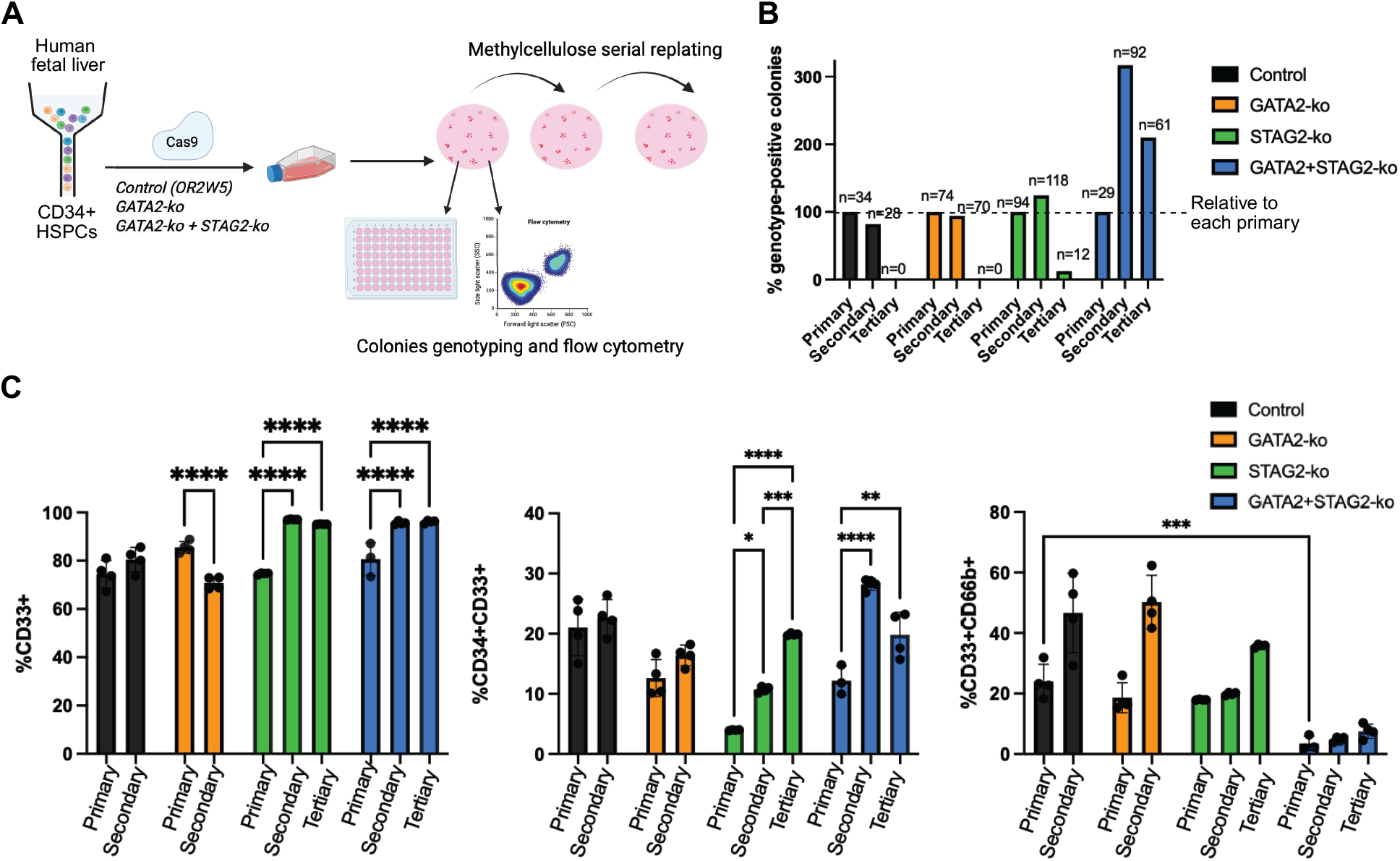
GATA2-ko + STAG2-ko confers higher HSC self-renewal *in vitro*. (A) Experimental overview of serial replating assay of control, GATA2-ko, STAG2-ko, and GATA2-ko + STAG2-ko-edited FL HSCs on methylcellulose colony forming media, with genotyping and immunophenotyping performed at each replating time point. (B) Percentage of genotype-positive colonies from each condition, including control, GATA2-ko, STAG2-ko, and GATA2-ko + STAG2-ko, after each replating (primary, secondary, and tertiary) relative to the primary plating, n = 29-74 genotype-positive colonies. (C) Immunophenotypic analysis of the output cells from each condition at each replating timepoint. CD33: myeloid lineage; CD66b: mature granulocyte marker; CD34: stem cell marker. *p < 0.05, **p < 0.01, ***p < 0.001, and ****p-value <0.0001, one-way ANOVA test, n = 3-4.

## Discussion

In this study, we explored the acquisition of a STAG2 knockout mutation in familial GATA2 deficient MDS, demonstrating its impact on increased HSC self-renewal and inflammation. Using CRISPR/Cas-9 mediated genome engineering, we induced GATA2 deficiency and STAG2-ko mutations in human HSPCs from prenatal FL. *In vivo*, xenotransplantation revealed a myeloid lineage bias induced by GATA2-ko + STAG2-ko mutations, a bias that was unique to xenografts with both mutations and not observed in xenografts with either GATA2-ko or STAG2-ko alone, indicating their synergistic effect.

While prior studies relying on genetically engineered mouse models have demonstrated myeloid-lineage skewing from both GATA2-ko^9,10^ and STAG2-ko^17^ alone, our study using a humanized model showed a required co-occurrence of the mutations to induce this pronounced myeloid bias. In fact, STAG2-ko is most common as a co-occurring mutation, rarely observed as an isolated mutation^29^. Our model, therefore, faithfully recapitulates this natural pattern of STAG2 as an acquired mutation, progressing a myeloid pre-disposition to high-risk.

Our inability to produce an MDS immunophenotype in our humanized model of GATA2 deficiency alone may be due to an insufficient length of time for engraftment. GATA2-deficient MDS is a disease that develops slowly over the course of childhood and adolescence and is usually not diagnosed until late teens or early twenties^10^, despite being a germline mutation. Therefore, the 12-week engraftment period of our GATA2-deficient HSCs in mice may not allow for the development of an observable phenotype. Furthermore, many individuals with germline GATA2 deficiency remain asymptomatic until the acquisition of additional somatic mutations^10^. Previous studies have also shown reduced fitness of GATA2 mutated HSPCs *in vivo*, both in mouse^9,10^ and humanized^11^ models, which could impact their ability to generate an MDS phenotype. Despite the lack of an MDS phenotype, we were able to demonstrate a pronounced loss of HSPC fitness transcriptionally in our model, showing a reduced number of primitive cell types in the GATA2-ko condition. A limitation of our study is that we were not able to separately analyze cells with heterozygous or homozygous GATA2 knockout, so immunophenotypic and single cell sequencing analyses are based on both homozygous and heterozygous loss of GATA2 function.

Beyond the immunophenotypic myeloid skewing observed in GATA2-ko + STAG2-ko xenografts, scRNA-seq revealed a marked expansion of HSCs and progenitors and identified these populations as the most transcriptionally perturbed relative to controls. In these primitive compartments, the double-mutant state was characterized by increased expression of stem cell maintenance programs alongside activation of inflammatory response pathways. Notably, pathways related to MYC target genes were downregulated in GATA2-ko + STAG2-ko HSCs. Although MYC is classically associated with proliferation^22^, it also plays a key role in balancing HSC self-renewal and differentiation, and reduced MYC activity may therefore favor stem cell maintenance at the expense of lineage commitment, providing a mechanistic explanation for the accumulation of primitive cells in this condition. In contrast, MYC signaling was upregulated in GATA2-ko HSCs, potentially explaining the absence of HSC expansion in the single-mutant setting. Together, the combination of enhanced stem cell maintenance and inflammatory signaling represents a transcriptional hallmark of MDS.

Within our scRNA-seq dataset, we identified ligand-target gene interactions between precursor to conventional dendritic cells (pre-cDCs) and HSCs in GATA2-ko + STAG2-ko xenografts, resulting in an upregulation of HLA-D genes in the HSC compartment. The upregulation of this MHC-II class gene often plays a role in decreasing HSC apoptosis^25^, thus contributing to the expansion of this cell type in our model. Pre-cDCs are often implicated in inflammatory signaling, including TNF-α and IFN-γ^30^. Therefore, this data suggests that the upregulation of inflammatory response pathways observed in HSCs in our model may be due to the production of ligands by more mature cells in the niche, such as dendritic cells. This inflammatory signaling could lead to an upregulation of MHC-II class genes, contributing to the expansion of the HSC compartment. Future studies could explore this inflammatory signaling as a potential therapeutic target in MDS.

The upregulation of MHC-II genes was also observed in the HSCs of GATA2-ko and STAG2-ko alone, albeit more prominently in STAG2-ko. This indicates the contribution of each gene mutation individually to the observed phenotype in the combined condition. Notably, HLA genes are often downregulated in full-blown leukemia as a tool of immune evasion^31^. Conversely, the upregulation of HLA-D genes here potentially underscores the preleukemic nature of our GATA2-ko + STAG2-ko MDS model.

Using *in vitro* assays, we were able to confirm the expansion of HSCs *in vitro* by demonstrating an increased colony-forming capacity of GATA2-ko + STAG2-ko HSCs over multiple weeks, indicating increased stem cell maintenance and self-renewal. We also were able to further validate the immature myeloid bias of GATA2-ko + STAG2-ko cells. The similar increase in colony-forming capacity and myeloid lineage bias in the STAG2-ko condition was expected, due to the observations from previous STAG2-ko mouse models^17^. Importantly, however, a combined GATA2 and STAG2 loss was required for a pronounced increase in colony-forming capacity in this assay. Finally, the decrease in colony-forming capacity over multiple weeks in the GATA2-ko condition is consistent with previous findings of the loss of self-renewal capacity in GATA2 deficient HSCs^11^.

Notably, despite an increase in HSC self-renewal upon STAG2-ko *in vitro*, we did not see an expansion of HSCs in the transcriptional data for STAG2-ko alone. This could be explained by a difference in experiment duration, with the colony data collected from a 4-week *in vitro* culture and the transcriptional data collected from a 12-week xenograft. One possibility is that STAG2-deficient HSCs exhibit a transient self-renewal advantage that is not sustained long term *in vivo*.

Collectively, our immunophenotyping, single-cell transcriptomic, and functional analyses demonstrate that combined GATA2 and STAG2 loss is required to induce an MDS-like state. While GATA2 deficiency alone was insufficient to initiate disease in our humanized system, STAG2 loss unmasked key features of high-risk progression, highlighting the cooperative nature of these lesions. Looking forward, this model provides a tractable platform for functional studies to define how inflammation remodels the HSC compartment in GATA2-ko + STAG2-ko MDS and to test whether inflammatory signaling creates therapeutic vulnerabilities that can be targeted.

## Acknowledgments

This work was supported by US National Institutes of Health (NIH) grants R01CA292503, R01CA290681 and R21CA301237, an Edward P. Evans Foundation Discovery Research Grant, by the HMTF Pilot Award Program for MDS Research, by an Alex’s Lemonade Stand Foundation Grant (22-25847), by a CureSearch for Children’s Cancer Grant, by a Blood Cancer United (formerly Leukemia and Lymphoma Society) Specialized Center of Research Program Grant (7039-25), by a V Foundation for Cancer Research Grant (V2024-015) and by a CURE Childhood Cancer Research Grant to E.W. In addition, E.W. is a Pew-Stewart Scholar supported by the Pew-Stewart Scholars Program for Cancer Research. M.Q.A. is supported by a grant from the Rally Foundation. This work was supported in part by the Bioinformatics for Next Generation Sequencing (BiNGS) shared resource facility of the Tisch Cancer Institute at the Icahn School of Medicine at Mount Sinai, which is partially supported by US NIH Cancer Center Support grant P30CA196521. This work was also supported in part through the computational and data resources and staff expertise provided by Scientific Computing and Data at the Icahn School of Medicine at Mount Sinai and supported by the Clinical and Translational Science Awards (CTSA) grant UL1TR004419 from the National Center for Advancing Translational Sciences. Research reported in this publication was also supported by the Office of Research Infrastructure of the NIH under award numbers S10OD026880 and S10OD030463.

We thank Zhihong Chen, Travis Dawson, Darwin D’Souza, Rachel Chen, Kai Nie and Seunghee Kim-Schulze from the Human Immune Monitoring Center (Mount Sinai) for single-cell library generation; Deniz Demircioglu from the Bioinformatics for Next Generation Sequencing (BiNGS) Shared Resource (Mount Sinai) for computational support; Edgardo Ariztia, Guillermo Villegas and Jordi Ochando from the Flow Cytometry CoRE (Mount Sinai) for assistance with flow cytometry; Franco Carlos, Lenny Martinez, Chineta Pullin, Kelly Yamada and Jonathan Cohen from the Center for Comparative Medicine and Surgery (Mount Sinai) for support with mouse work; We thank the Developmental Origin of Health and Disease (DOHaD) Biorepository, including Ya-Wen Chen, Mikal Kizilbash, and Meghana Sreenath. We thank members of the Wagenblast lab for their comments on the manuscript. Figures were created in BioRender.

## Author Contributions

G.F., M.Q.A., and E.W. designed, performed, analyzed experiments, and prepared figures. L.L., N.P., S.C., S.M. and D.H. provided computational expertise, analyzed all scRNA-seq and GoT-ChA datasets, and prepared figures. S.A., P.F., K.E., I.G.M., M.Z., S.S. and C.C. assisted with *in vivo* experiments. L.M. assisted with GoT-ChA experiments. Z.D. assisted with sample processing. G.F. wrote the manuscript with assistance from E.W. E.W. supervised the study.

## Supplementary Figure Legends

**Supplementary Figure 1. Characterization of the xenograft immunophenotype.** (A) UMAP showing GATA2 homozygous mutant, heterozygous mutant, and wild-type cells in the GoT-ChA dataset from GATA2-ko xenograft. (B) Table showing the number of cells within each genotype classification and their respective percentages out of the total number of genotyped cells. HET is heterozygously mutated for GATA2, HOM is homozygously mutated for GATA2, WT is wild-type and NA is unknown genotype. (C) Heat map showing transcription factor motif enrichments across cell types in our GoT-ChA dataset (D) Immunophenotypic analysis from the non-injected bone marrow of control, GATA2-ko, STAG2-ko, and GATA2-ko + STAG2-ko primary xenografts. CD45: overall human cell engraftment; CD33: myeloid lineage; CD19: lymphoid lineage; CD71: immature erythroid marker; GlyA: mature erythroid marker. The line represents the mean value. **p-value <0.01, Kruskal-Wallis Test, n = 5-18 mice each.

**Supplementary Figure 2. Transcriptional landscape of different xenograft conditions.** (A) UMAPs representing xenografts of control, GATA2-ko, STAG2-ko, and GATA2-ko + STAG2-ko with cell-type annotations. Total cell number for each condition: Control: 23,398 cells, GATA2-ko: 8,466 cells, STAG2-ko: 6,458 cells and GATA2-ko + STAG2-ko: 6,402 cells. (B-C) Result from Augur bioinformatics analysis showing prioritization of cell types based on their molecular response. Showing comparison of GATA2-ko versus matched control and (C) STAG2-ko versus matched control. (D) Result from Augur analysis showing significantly differentially prioritized cell types between GATA2-ko and GATA2-ko + STAG2-ko.

**Supplementary Figure 3. Validation of distinct transcriptional landscape of GATA2-ko + STAG2-ko progenitor cells.** (A-D) Over Representation Analysis (ORA) on differentially expressed (A) cycling progenitor genes between GATA2-ko + STAG2-ko versus control, (B) early GMP genes between GATA2-ko + STAG2-ko versus control, HSC genes between GATA2-ko versus control, (D) HSC genes between STAG2-ko versus control, and (E) HSC genes between GATA2-ko + STAG2-ko versus GATA2-ko. Enriched pathways and biological processes were identified based on differentially expressed genes. Significant hallmark pathways are highlighted in bold and with an asterisk (*) based on p-value < 0.05. (E-F) Volcano plot of differentially expressed genes between (F) GATA2-ko + STAG2-ko and control in the cycling progenitor compartment and (G) GATA2-ko + STAG2-ko and control in the early GMP compartment. Genes associated with inflammation are marked in green and genes associated with self-renewal are marked in blue. All genes with significant adjusted p-values < 0.05 and log2FC > 1 are colored.

