## Supplemental Methods for "A CRISPR-Based Humanized Model Reveals Cooperative Role of STAG2 Loss in Familial GATA2-Deficient MDS Progression"

### Method Details

Fetal liver CD34+ HSPC isolation

The fetal liver sample was finely minced using razor blades and was then put into two 50 ml Falcon tubes, each with 40 ml of pre-warmed IMDM medium (Thermo Fisher), 5 ml of collagenase IV (Stem Cell Technologies), and 25 µl of 10 mg/ml DNAse I (Roche). The 50 ml tubes were incubated on a shaker at 37° C for 30 min. Following incubation, the dissociated sample was filtered through a 40 µm cell strainer (Thermo Fisher). The remaining tissue pieces were pushed through the strainer using the black rubber end of a 5 ml syringe (BD). The cells were then centrifuged at 350 x g for 10 min. The red blood cells were lysed with 5 ml ammonium chloride (StemCell Technologies) per 50 ml tube for 5 min at room temperature. Lysis was stopped with IMDM medium (Thermo Fisher), and cells were centrifuged at 350 x g for 10 min. The remaining cell pellets were combined and re-suspended in 500 µl MACS auto-running buffer (Miltenyi Biotec). Subsequently, CD34+ cells were subsequently enriched using the human CD34 MicroBead kit (Miltenyi Biotec) according to the manufacturer's protocol. In total, 12 LS columns (Miltenyi Biotec) were used per fetal liver sample. CD34+ fetal liver cells were viably stored in a cryopreservation solution consisting of 90 % FBS (Cytiva) and 10 % DMSO (Sigma) in liquid nitrogen.

Primary CD34+ HSPCs

CD34+ enriched HSPCs were thawed via slow dropwise addition of thawing medium, made up of X-VIVO 10 medium (Lonza) with 50 % FBS (Cytiva) and DNase I (100 µg/ml, Roche). Cells were centrifuged at 350 x g for 10 min and then re-suspended in PBS (Cytiva) + 2.5 % FBS (Cytiva). CD34+ HSPCs were resuspended in 1000 µl PBS + 2.5 % FBS per 5×x$10^6$ cells and stained for 60 min at 4° C. The following antibodies were used (volume per 1×x$10^6$ cells): CD45 V500 (1:100, BD) and CD34 APC-Cy7 (1:200, BD). Afterwards, cells were washed and resuspended. The viability stain 450 (BD) was added (1:1000). CD34+ HSPCs were sorted using the FACSMelody (BD). Cell sorting purity checks were performed after each sorted sample (>95 %).

CD34+ HSPCs *in vitro* culture

FACS-sorted CD34+ HSPCs were cultured in serum-free X-VIVO 10 medium (Lonza) supplemented with 1 % bovine serum albumin fraction V (Roche), 1x L-glutamine (Thermo Fisher), 1×x penicillin-streptomycin (Thermo Fisher) and the following cytokines (Biotechne): FLT3 Ligand (100 ng/mL), G-CSF (10 ng/mL), SCF (100 ng/mL), TPO (15 ng/mL) and IL-6 (10 ng/mL). Cells were cultured in 96 well round-bottom plates (VWR).

CRISPR/Cas9 RNP electroporation

Sorted CD34+ HSPCs were cultured for 48 hours in serum-free X-VIVO 10 medium (Lonza) as previously described[32]. CRISPR/Cas9 RNP electroporations were performed using chemically synthesized gRNAs (IDT), recombinant Cas9 nuclease (IDT), and the 4D-Nucleofector (Lonza). For the preparation of the RNP complex, Alt-R CRISPR/Cas9 crRNA and tracrRNA (IDT) were reconstituted to a concentration of 200 µM in TE Buffer (IDT). The crRNA and tracrRNA were then mixed in a 1:1 ratio and annealed using a thermocycler at 95° C for 5 min, followed by cooling to room temperature. For each electroporation reaction, 1.2 µl of crRNA:tracrRNA complex, 1.7 µl of Cas9 protein, and 2.1 µl of PBS (Cytiva) were combined in a low-binding Eppendorf tube and incubated at room temperature for 15 min to form the complex. If multiple genes were targeted in the same electroporation reaction, such as for GATA2-ko + STAG2-ko, 1.2 µl of each crRNA:tracrRNA complex, 1.7 µl of Cas9 protein, and 0.9 µl of PBS were combined into the same Eppendorf tube. After incubation at room temperature for 10 min, 1 µl of 100 µM electroporation enhancer (IDT) was added to the mixture.

Pre-cultured cells were washed with pre-warmed PBS and centrifuged at 350 x g for 10 min. Approximately $2.5 \times x10^5$ cells were resuspended in 20 µl of Buffer P3 (Lonza) per reaction and added to the tube containing the CRISPR/Cas9 gRNA RNP complex. The mixture was thoroughly mixed by pipetting up and down and then transferred to the electroporation chamber (Lonza).

The cells were electroporated using the 4D-Nucleofector with the program DZ-100. Immediately following electroporation, 180 µl of pre-warmed X-VIVO 10 medium (as described above) was added to the electroporation chamber, and cells were transferred to a 96-well round-bottom plate. The electroporated cells were allowed to recover overnight at 37° C in a tissue culture incubator. For each CRISPR/Cas9 electroporation, a small subset of cells was cultured for 5-7 days to validate CRISPR/Cas9 efficiency.

<u>gRNA sequences</u>

The gRNA sequence for GATA2 was designed using Benchling based on naturally occurring mutation sites in the second zinc-finger domain[8]. The gRNA sequence for STAG2 was designed and tested in a previous study[18]. For control, two gRNAs targeting the olfactory receptor OR2W5 were used.

<u>Methylcellulose colony formation assay</u>

For methylcellulose colony formation assays, CD34+ HSPCs sorted from human fetal liver, were cultured in X-VIVO-10 medium (as described above). After 48 hours of culture, the cells proceeded to CRISPR/Cas9 RNP electroporation as described. Two days post-electroporation, cells were collected and counted, and a specific number of cells were resuspended in MethoCult H4034 optimum methylcellulose medium (StemCell Technologies) and plated in 35 mm cell culture dishes. These cells were labeled as day 1 cells. After 12-14 days, individual colonies were picked and genotyped to analyze the CRISPR/Cas9-edited loci. Colonies with editing efficiency >75% were defined as homozygous and colonies with editing efficiency <75% were defined as heterozygous. In the case of the serially replating assay, to further analyze the immunophenotypes of the colonies derived from methylcellulose assays, remaining cells were collected and pooled by adding 3 mL of PBS into each 35 mm dish and washed three times with PBS with 2.5% FBS. Approximately 100,000 of the collected cells were then stained with antibodies, and their immunophenotypes were analyzed by flow cytometry. The remaining cells were counted and replated on methylcellulose medium in the same method as previously described. In the case of the inflammation challenge assay, cells were maintained in liquid culture, with media changed or cells split every other day to ensure high viability. At subsequent time points, including day 8 and day 15 cells were plated again on methylcellulose medium as described for day 1. 20-40 individual colonies per condition were picked, washed with PBS, and stored in a 96-well PCR plate (Eppendorf) at -80° C. For genotyping, genomic DNA isolation and PCR amplification were performed. Genomic DNA was isolated from colonies using the Agencourt GenFind V3 kit (Beckman Coulter). The isolation process was performed in 96-well PCR plates (Eppendorf) using a magnetic stand (Thermo Fisher), following the manufacturer's protocol with the following volume modifications: 100 µl lysis buffer, 2.3 µl of proteinase K, 75 µl magnetic particles, 200 µl wash buffer 1, 125 µl wash buffer 2 and 60 µl TE buffer (IDT) for elution. PCR was used to amplify the CRISPR/Cas9-modified genomic locus. Each PCR reaction contained 23 µl of genomic DNA, 1 µl of forward and reverse primer (10 µM, IDT), and 25 µl of AmpliTaq Gold 360 master mix (Thermo Fisher). The PCR program was: 95° C for 10 min, followed by 95° C for 30 s, 56° C for 30 s and 72° C for 1 min (40 cycles), and then 72° C for 7 min. For control-(OR2W5), expected deletions were confirmed through gel electrophoresis of the PCR products. For GATA2-ko and STAG2-ko, PCR products were purified, and Sanger sequencing was performed using the reverse PCR primer. Chromatographs were analyzed using TIDE[33] to verify editing efficiencies.

### Primary xenotransplantation

The electroporated cells were allowed to recover overnight at 37° C in a tissue culture incubator. Then, cells were transplanted into sublethally irradiated NSG mice via intrafemoral injections. Mice were anesthetized with isoflurane, and their left knee was secured in a bent position. A 27-gauge needle was used to drill a hole into the left femur, followed by the injection of CRISPR/Cas9-edited HSPCs in 30 µl of PBS using a 28-gauge ½ cc syringe (BD). Mice from the primary xenotransplantation of CRISPR/Cas9-edited HSPCs were euthanized at week 12. Engraftment was considered positive if the human CD45+ engraftment level in the bone marrow was greater than 1%, with the population presenting as dense clusters rather than dispersed signals.

### Genotyping of xenotransplantations

To assess the efficiency of each xenograft, genomic DNA was isolated from bulk bone marrow cells using the Agencourt GenFind V3 kit (Beckman Coulter), following the same procedure as previously described. PCR and resulting analyses were performed using the same protocol as previously described for assessing editing efficiencies.

### Flow cytometry

For xenografted cells, the bone marrow, including left and right femurs, of each NSG mouse was flushed with 1 ml staining buffer (PBS + 2.5 % FBS). Collected cells were centrifuged at 350 x g for 10 min and resuspended in 500 µl staining buffer. For antibody staining, 50 ul of cells per sample were mixed with 50 µl of antibody mix and incubated for 60 minutes at 4° C, protected from light. After staining, cells were washed once with staining buffer and resuspended in 250 µl staining buffer. In addition, 25 µl of each sample were transferred to a 96-well PCR plate (Eppendorf) and stored at -80°C for genotyping. The remaining cells were used for flow cytometry analysis. The following antibodies were used (all from BD, unless stated otherwise): CD45 V500 (1:100), CD33 BV711 (1:100), CD19 PE-Cy7 (1:100), CD41 PE-Cy5 (1:200, Beckman Coulter), GlyA PE (1:100, Beckman Coulter), CD71 BV650 (1:100), CD3 FITC (1:100), CD34 APC-Cy7 (1:100), CD117 APC (1:100), CD66b FITC (1:100), CD14 APC (1:100). A pan-lineage panel was designed and used for primary xenograft analysis including: CD45, CD19, CD3, CD33, CD41, GlyA, CD71, CD34 and CD117. A myeloid panel was designed and used for serial methylcellulose replating analysis including CD45, GlyA, CD33, CD66b, CD14, CD34, CD41, and CD117. Flow cytometry was performed using the FACSFortessa (BD Biosciences). The remaining unstained cells from each xenografted mouse were viably frozen and stored in liquid nitrogen. For *in vitro* cultured cells, at least 100,000 cells were collected and stained for each experiment. The cells were stained for 60 minutes at 4° C, protected from light. Cells were then washed once with staining buffer and resuspended in 250 µl of staining buffer. Flow cytometry analysis was performed using the FACSFortessa (BD) or Attune (Thermo Fisher). All data were analyzed with FlowJo.

### ScRNA-seq library generation

Xenografted cells were thawed following the previously described protocol. Human cells were isolated using mouse depletion kit (Miltenyi Biotec), then cells were stained with CD45 V500 (1:100), CD34 APC-Cy7 (1:100), and CD19 PE-Cy7 (1:100), in staining buffer and incubated for 60 minutes at 4° C, protected from light. Fixable Viability Stain 450 (1:1000) was added 10 minutes before sorting and viable human CD45+ cells were sorted using FACSMelody (BD). On average, 1-2x$10^5$ cells per sample was submitted for scRNA-seq and scATAC-seq library generation.

Viability of single cells was assessed using Acridine Orange/Propidium Iodide viability staining reagent (Nexcelom), and debris-free suspensions of >80% viability were deemed suitable for experiments. The Chromium 10x Genomics 5' GEX v2 protocol was used for scRNA-seq. Briefly, Gel-Bead in Emulsions (GEMs) were generated on the sample chip using the Chromium X system. Barcoded cDNA was extracted from the GEMs following post-GEM RT cleanup and amplified for 12 cycles. For GEX library preparation, amplified cDNA was fragmented and subjected to end-repair, poly A-tailing, adapter ligation, and 10x-specific sample indexing, following the manufacturer's protocol. The GEX libraries were quantified using TapeStation (Agilent) and QuBit (ThermoFisher) and sequenced in paired-end mode on a NovaSeq 6000 instrument (Illumina), targeting a depth of 25,000 reads per cell. Raw Fastq files were aligned to the GRCh38 reference genome (2020-A) using Cell Ranger v5.0.1 (10x Genomics).

### ScRNA-seq data quality control and pre-processing

Sequenced FASTQ files were aligned, filtered, barcoded, and unique molecular identifier (UMI) counted using Cell Ranger Chromium Single Cell RNA-seq (v 7.1.0 and v5.0.1)

(10X Genomics) with the hg38 human genome reference (version 2020-A; RRID: SCR_017344). Cells were filtered to retain those with ≥1000 UMIs, ≥400 expressed genes, and <5% of reads mapping to the mitochondrial genome. UMI counts were then normalized to a total of 10,000 UMIs per cell and log-transformed with a pseudocount of 1 using the “LogNormalize” function in the Seurat package (version 4.0.3; RRID: SCR_016341)[34,35]. The top 2,000 highly variable genes were identified with the “vst” method in the “FindVariableFeatures” function, and the data were scaled using the “ScaleData” function. Cell cycle scoring was performed using the “CellCycleScoring” function, which assigns cell cycle phase scores (G1, S, and G2/M) based on canonical marker gene expression.

#### ScRNA-seq data dimensionality reduction and integration

Principal component analysis was performed using the top 2,000 highly variable features (“RunPCA” function) and the top 30 principal components were used in the downstream analysis. K-Nearest Neighbor graphs were obtained using the “FindNeighbors” function whereas the UMAPs were obtained using the “RunUMAP” function. The Louvain algorithm was used to cluster cells based on expression similarity. The top cluster biomarkers were identified by “FindAllMarkers” function.

#### ScRNA-seq data cell type annotation and differential biomarker analysis

To annotate clusters with relevant cell types, the sc-transcriptomes were projected against a hematopoietic reference map using BoneMarrowMap R package[17=8]. To visualize the contribution of each annotated cell type across genotypes, the percentage of cells in each cell type within each genotype was represented with bar plots. Differential markers for each cluster were identified using the Wilcox test (“FindMarkers” function) with adjusted p-value < 0.01, absolute log2 fold change > 0.25, and minimum 10 % of cells expressing the gene in both comparison groups using 1,000 random cells to represent each cluster. The differentially expressed genes were then used to perform overrepresentation analysis (ORA) with enricher function from the clusterProfiler package using the collections from the Molecular Signatures Database (MSigDB). The enriched pathways were visualized using barplots.

#### ScRNA-seq data cell type prioritization and cell-cell communication

We used Augur[21] to quantify the transcriptional responsiveness of each cell type to the knockout perturbation. For each comparison, the dataset was subset to include only the knockout and its matched control. Augur estimates the separability of experimental conditions within each cell type using machine-learning-based classification, generating an area-under-the-curve (AUC) score as a measure of perturbation responsiveness. Raw count matrices and corresponding cell annotations were provided as input, and sample-level condition labels were used to define the perturbation. All analyses were performed using default Augur parameters. Higher AUC values indicate greater transcriptomic separability between conditions and thus greater perturbation sensitivity.

Cell-cell communication analysis was performed using NicheNet[26], following the standard workflow and default parameters. We focused on hematopoietic stem cells (HSCs) as the receiver population and defined expressed genes in HSCs to identify active receptors and downstream target genes. A curated set of non-lymphoid sender populations was used by removing all lymphoid cell types prior to defining expressed ligands. Potential ligands were restricted to those whose cognate receptors were expressed in HSCs. Differentially expressed genes in HSCs between the knockout and control conditions were used as the gene set of interest, with all expressed HSC genes serving as the background. Ligand activity was inferred using NicheNet's ligand-target matrix to compute AUPR-based ligand activity scores.

<u>Genotyping of targeted loci with chromatin accessibility (GoT-ChA) library preparation</u>

Xenografted cells were thawed following the previously described protocol. Human cells were isolated using mouse depletion kit (Miltenyi Biotec), then cells were stained with CD45 V500 (1:100), CD34 APC-Cy7 (1:100), and CD19 PE-Cy7 (1:100), in staining buffer and incubated for 60 minutes at 4° C, protected from light. Fixable Viability Stain 450 (1:1000) was added 10 minutes before sorting and viable human CD45+ cells were sorted using FACSMelody (BD). Cells were subjected to GoT-ChA[19]. Briefly, the nuclei isolation procedure is the same as scATAC-seq, and the isolated nuclei were processed according to the Chromium Next GEM Single Cell ATAC Solution User Guide (version CG000209 Rev F, 10×x Genomics) with the following modifications:

a. GEM Generation and Barcoding: During the GEM generation and barcoding reaction (step 2.1 in 10x protocol), 1 µl of a 22.5 µM GoT-ChA primer mix was added to the barcoding mixture. The primers used in this step were: GATA2-ko forward: 5'-TCGTCGGCAGCGTCAGATGTGTATAAGAGACAGGCACTGAAGGGGGATGAC*T*T*C

Reverse: GCCCTCTGAAAACTGGTG*G*T*T

b. Post-GEM Incubation and Cleanup: During the post-GEM incubation clean-up (step 3.2 in 10x protocol), 45.5 µl of Elution Solution I was used to elute material from SPRIselect beads (Beckman Coulter). A total of 5 µl was used for GoT-ChA library construction, and the remaining 40 µl was used for ATAC library construction as per the standard protocol.

c. GoT-ChA Library Construction: Two additional PCR reactions were performed using the 5 µl set aside during step 3.2 to generate the GoT-ChA library. The first PCR amplifies genotyping fragments before sample indexing. The primers used were: P5 (binds to the P5 Illumina sequencing handle): 5'-GATACGGCGACCACCGAGATCTACAC. GoT-ChA nested (biotinylated nested primer with a partial TruSeq small RNA read 2 handles): 5'-/5Bioag/ CCTTGGCACCCGAGAATTCCATGCCTCTAGGTTAACAGGCCA.
Thermocycling conditions are 95° C for 3 minutes, followed by 15 cycles of 95° C for 20 s, 65° C for 30 s, and 72° C for 20 s. The final extension is 72° C for 5 minutes, followed by holding at 4° C. After a 1.2 x SPRIselect cleanup, biotinylated PCR products were bound using Dynabeads M-280 Streptavidin magnetic beads (Thermo Fisher) at room temperature for 15 minutes. The beads were washed

twice with 1×x SSPE buffer and once with 10 mM Tris-HCl (pH 8.0) before resuspending in water. The bead-bound fragments were amplified and indexed using the following primers: P5 (for Illumina sequencing handle), RPI-X (adds a sample index and P7 Illumina sequencing handle: CAAGCAGAAGACGGCATACGAGATXXXXXXXXGTGACTGGAGTTCCTTGGC ACCCGAGAA TTCCA, where X represents a user-defined sample index). Thermocycling conditions are 95° C for 3 minutes, followed by 6-10 cycles of 95° C for 20 s, 65° C for 30 s, and 72° C for 20 s, final extension: 72° C for 5 minutes, followed by holding at 4° C. The final libraries were quantified using a High-Sensitivity DNA Chip (Agilent Technologies) run on a Bioanalyzer 2100 system (Agilent Technologies). Libraries were sequenced on the NovaSeq 6000 system (Illumina) with the following parameters: 50/8/16/50 for ATAC and 50/16/50 for the GoT-ChA library. The sequencing depth is 25,000 read pairs per nucleus for ATAC libraries and 5,000 read pairs per nucleus for GoT-ChA libraries.

GoT-ChA data analysis

For GoT-ChA, the FASTQ files from chromatin accessibility data were processed to generate the cell-by-gene count matrix with CellRanger v.5.0.1 using default parameters, with GRCh38 as the human genome reference. The data were filtered to retain cells with at least 3,000 peaks detected, a minimum of 50% of reads in peaks, nucleosome signal below 4 and TSS enrichment score above 3. Dimensionality reduction was performed in Seurat (v.5.1.0) applying term frequency-inverse document frequency (TF-IDF) followed by single value decomposition for latent semantic indexing (LSI). For graphical representation, dimensions were further reduced via UMAP, removing the first LSI dimension which showed a high correlation (> 0.5) with sequencing depth. Cell types were assigned based on gene activity scores as calculated by the GeneActivity function from Seurat (v.5.1.0) and concordance with the labels assigned via bridge dictionary mapping[35]. Transcription factor motif accessibility was calculated via chromVar[36]. Genotyping FASTQ files were processed with the Gotcha R package (v.1.0), using wild-type and GATA2-ko sequences for perfect match. Genotype calls were included in Seurat object metadata for downstream analysis. Differential analysis was performed via differential linear mixture models using the DiffLMM function from Gotcha (v.1.0) as previously done[19].

Statistical analysis

Statistical analysis was performed using GraphPad Prism software Version 10.2.3. Statistical significance was calculated using a Kruskal-Wallis test, unless otherwise noted. This non-parametric test was chosen because it is useful for comparing multiple conditions with a large range of sample sizes. Other relevant statistical methods are detailed in the methods section and figure legends. For all analyses, $p < 0.05$ was considered statistically significant.
