## Supplementary Figure 1 for "A CRISPR-Based Humanized Model Reveals Cooperative Role of STAG2 Loss in Familial GATA2-Deficient MDS Progression"

A

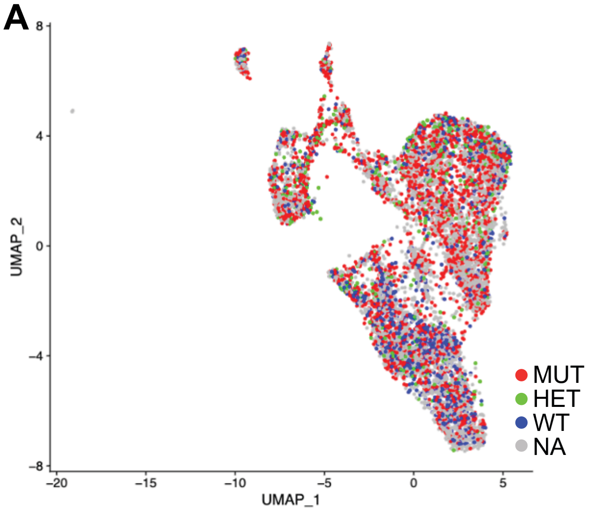


B

| Genotype | Number of cells | Percent of genotyped cells |
|---|---|---|
| HET | 511 | 12% |
| HOMO | 2757 | 63% |
| WT | 1087 | 25% |
| NA | 4055 | N/A |

C

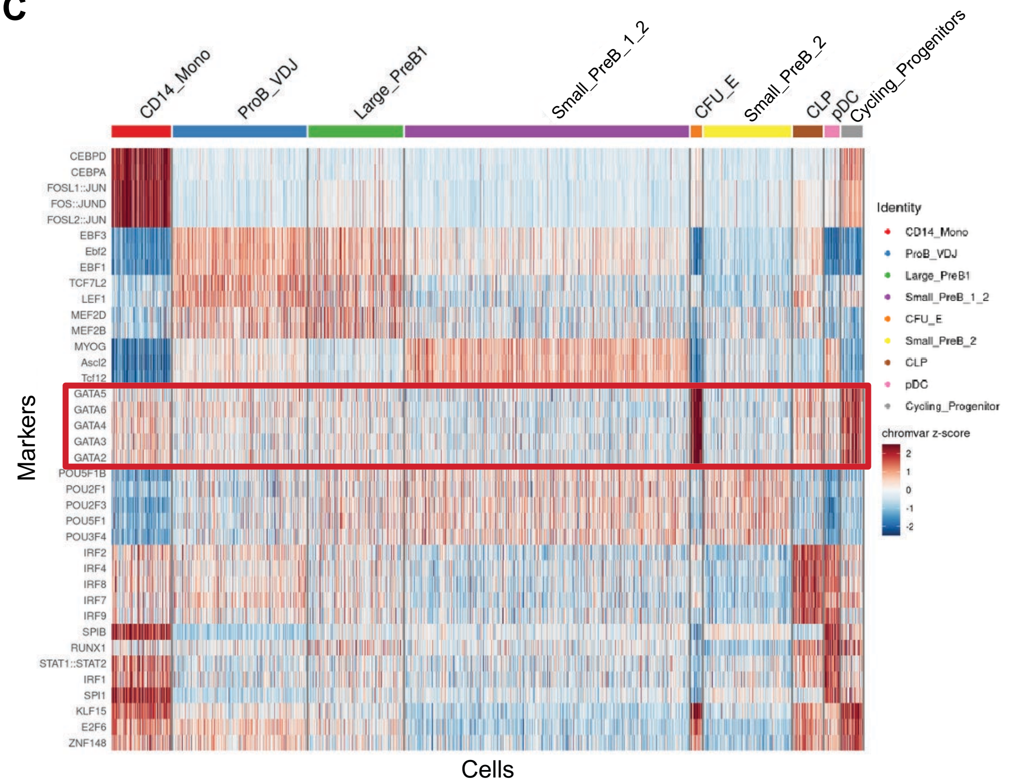


D

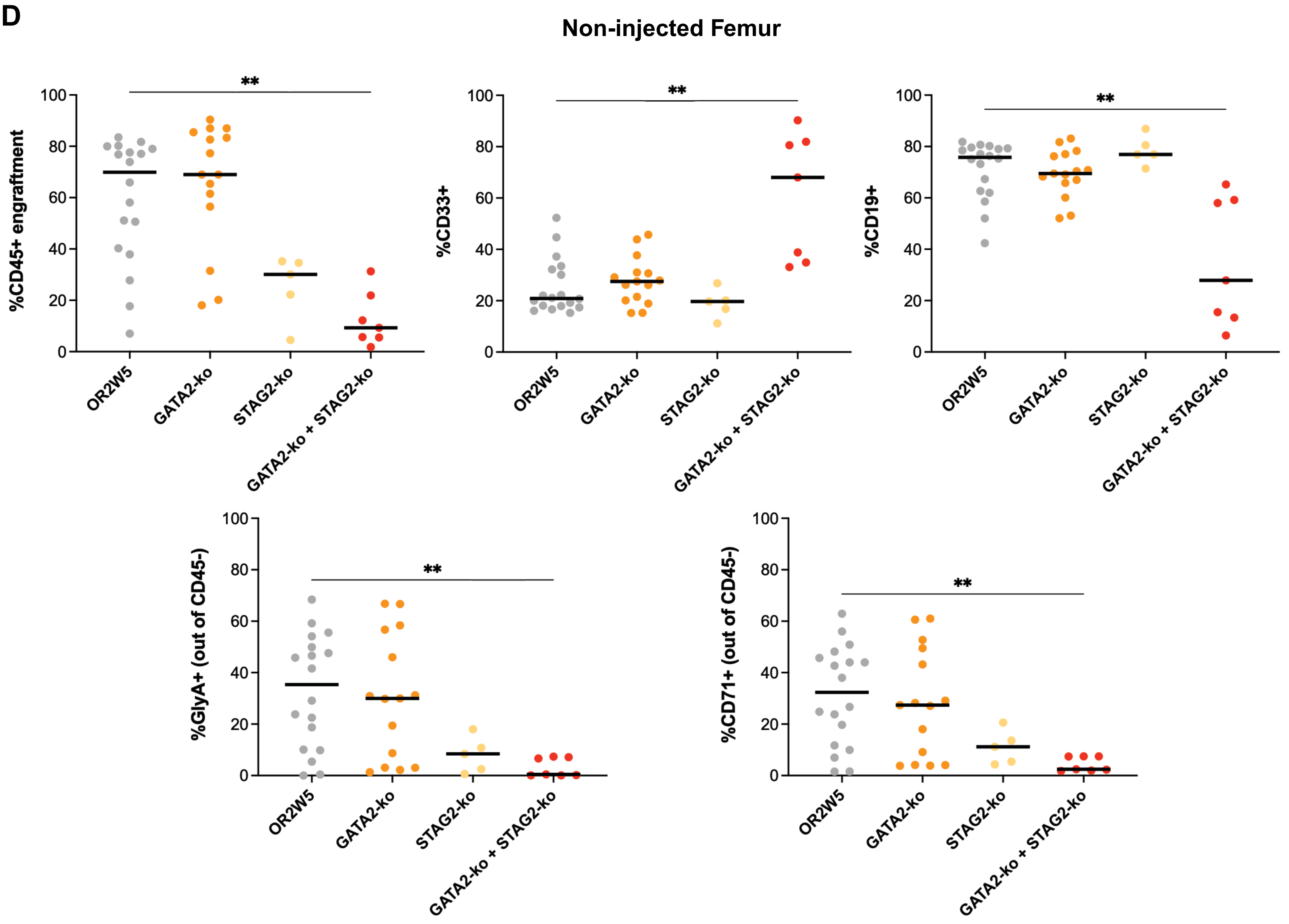
