## Supplementary figures and images for "A CRISPR-Based Humanized Model Reveals Cooperative Role of STAG2 Loss in Familial GATA2-Deficient MDS Progression"

### Supplementary Figure 2

##### Supplementary Figure 2

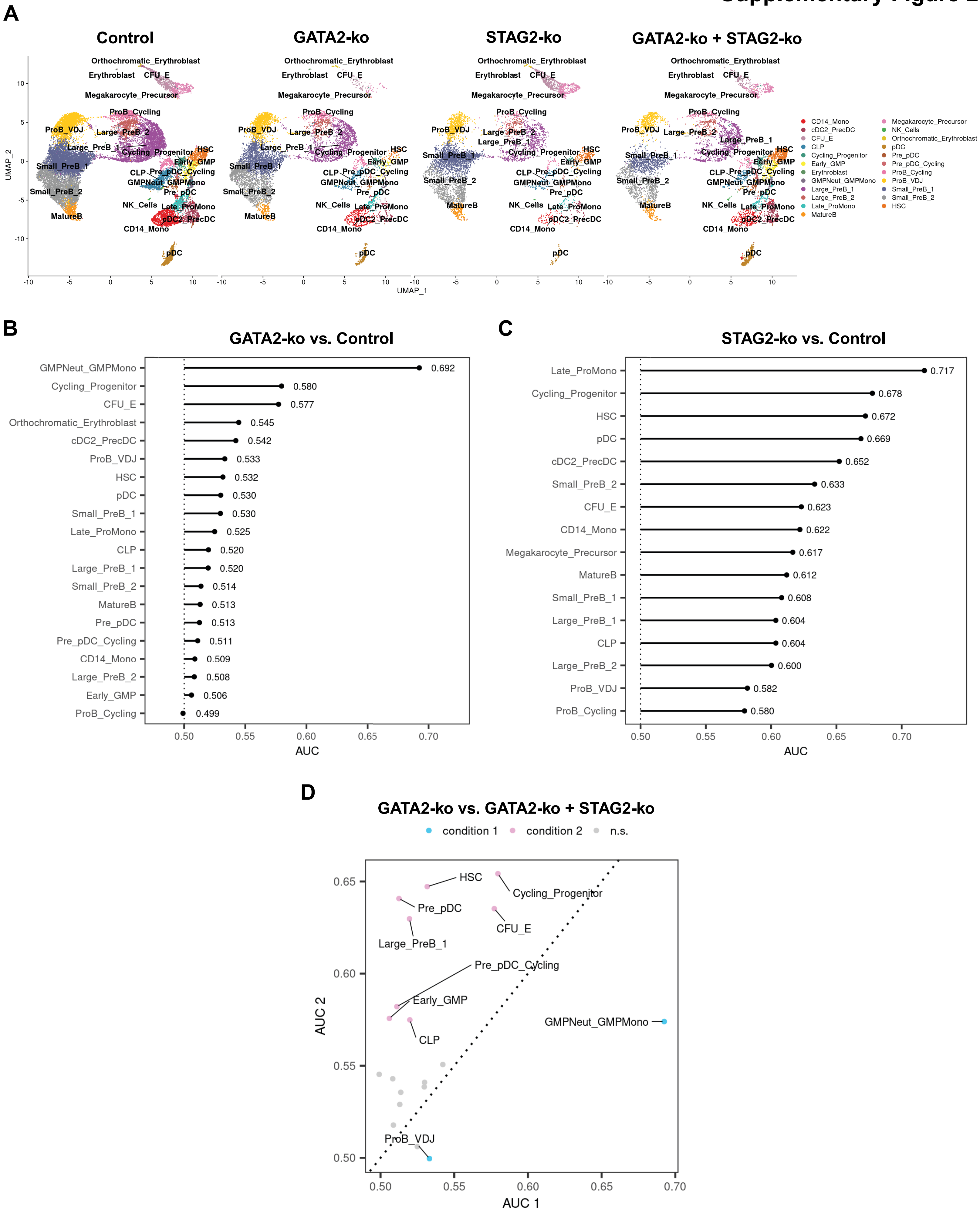

### Supplementary Figure 3

#### Supplementary Figure 3

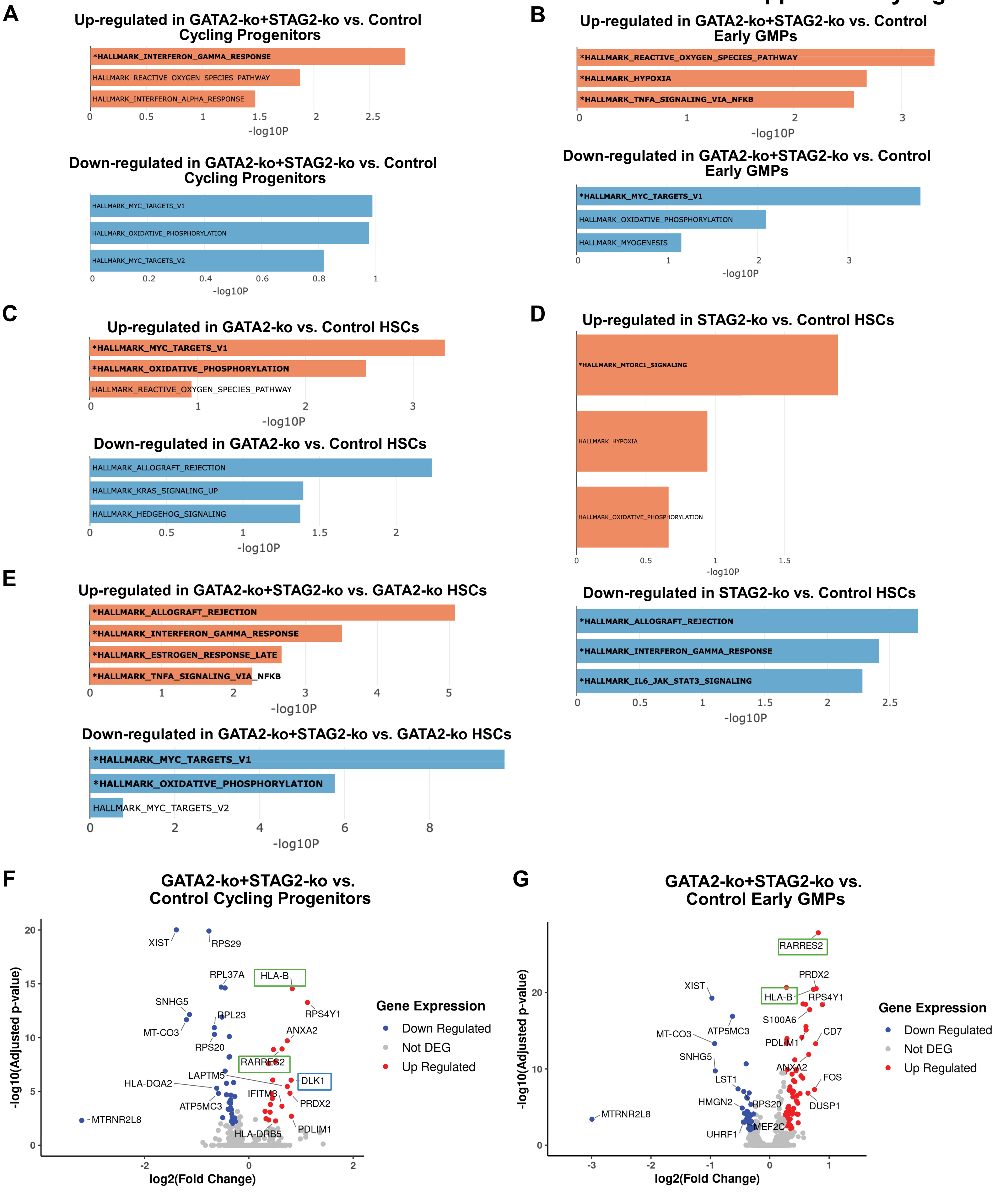
